# Wilson Statistics: Derivation, Generalization, and Applications to Electron Cryomicroscopy

**DOI:** 10.1101/2021.05.14.444177

**Authors:** Amit Singer

## Abstract

The power spectrum of proteins at high frequencies is remarkably well described by the flat Wilson statistics. Wilson statistics therefore plays a significant role in X-ray crystallography and more recently in electron cryomicroscopy (cryo-EM). Specifically, modern computational methods for three-dimensional map sharpening and atomic modelling of macromolecules by single particle cryo-EM are based on Wilson statistics. Here we provide the first rigorous mathematical derivation of Wilson statistics. The derivation pinpoints the regime of validity of Wilson statistics in terms of the size of the macromolecule. Moreover, the analysis naturally leads to generalizations of the statistics to covariance and higher order spectra. These in turn provide theoretical foundation for assumptions underlying the widespread Bayesian inference framework for three-dimensional refinement and for explaining the limitations of autocorrelation based methods in cryo-EM.

## 1 Introduction

The power spectrum of proteins is often modelled by the Guinier law at low frequencies and the Wilson statistics at high frequencies. At low frequencies, there is a quadratic decay of the power spectrum characterized by the moment of inertia of the molecule (e.g., its radius of gyration). At high frequencies, the power spectrum is approximately flat. In structural biology, it is customary to plot the logarithm of the spherically-averaged power spectrum of a three-dimensional structure as a function of the squared spatial frequency. This Guinier plot typically depicts the two different frequency regimes. It is not surprising that these laws are of critical importance in structural biology, with applications in X-ray crystallography [1] and cryo-EM [2]. However, while Guinier law has a very simple mathematical derivation based on a Taylor expansion, in the literature we could only find heuristic arguments in support of Wilson statistics, such as the original argument provided by Wilson in his seminal 1-page Nature paper [3]. Here we provide a rigorous mathematical derivation of Wilson statistics in the form of Theorem 3 and derive other forms of statistics with potential application to cryo-EM. The main ingredients to our analysis are a scaling argument, basic probability theory, and modern results in Fourier analysis that have found various applications within mathematics (such as the distribution of lattice points in domains), but their application to structural biology appears to be new.

### 1.1 Random bag of atoms

The model underlying Wilson statistics is a random “bag of atoms”, where the random “protein” consists of *N* atoms whose locations *X*_1_,*X*_2_,…, *X_n_* are independent and identically distributed (i.i.d). For example, each *X_i_* could be uniformly distributed inside a container 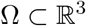 such as a cube or a ball, though other shapes and non-uniform distributions are also possible. The electron scattering potential 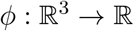 of the protein is modelled as

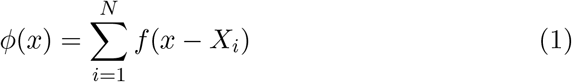

where *f* is a bump function such as a Gaussian, or a delta function in the limit of an ideal point mass. For simplicity of exposition, we assume that the atoms are identical. Otherwise, one can use different *f*’s to describe the scattering from each atom type. The Fourier transform of (1) is given by

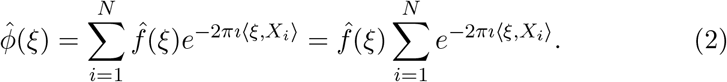

### 1.2 Wilson statistics

Wilson’s original argument [3] uses (2) to evaluate the power spectrum as follows

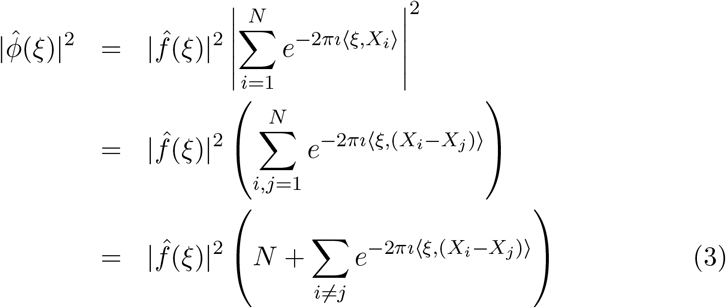

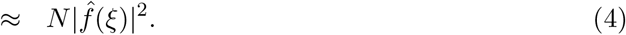

Wilson argued that the sum of the complex exponentials in (3) is negligible compared to *N*, as those terms wildly oscillate and cancel each other, especially for high frequency ξ. We shall make this hand wavy argument more rigorous and the term “high frequency” mathematically precise. Note that for an ideal point mass 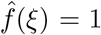, and (4) implies that the power spectrum is flat, i.e., 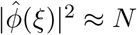.

The challenge is to show that there is so much cancellation that adding *O*(*N*^2^) oscillating terms of size *O*(1) in (3) is negligible compared to *N*. For a random walk, the sum of *O*(*N*^2^) i.i.d zero-mean random variables of variance *O*(1) is *O*(*N*) (the square-root of the number of terms). In order to show that the sum is negligible compared to *N*, additional cancellation must be happening. The role that ξ plays also needs to be carefully analyzed, as for ξ = 0, clearly 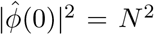. What is the mechanism by which 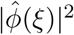 decays from *N*^2^ to *N* as ξ increases?

## 2 Derivation of Wilson Statistics

### 2.1 *N*^1/3^ scaling

Since *X*_1_,…,*X_N_* are i.i.d, one might be tempted to apply the Central Limit Theorem (CLT) to (2) and conclude that 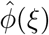 is approximately a Gaussian, for which the mean and variance can be readily calculated as done in [4]. However, one should proceed with caution, because if the container Ω is fixed, then in the limit *N* → ∞, the density of the atoms also grows indefinitely, whereas the density of atoms in a protein is clearly bounded. If the density of the atoms is to be kept fixed, the container Ω has to grow with *N*. To make this dependency explicit, we denote the container by Ω_*N*_. The volume of the container Ω_*N*_ must be proportional to *N*. The length-scale is therefore proportional to *N*^1/3^, that is, Ω_*N*_ = *N*^1/3^Ω_1_, or *X_i_* = *N*^1/3^*Y_i_* with *Y_i_* ~ *U*(Ω_1_) in the uniform case, and i.i.d in general. The Fourier transform (2) is rewritten as

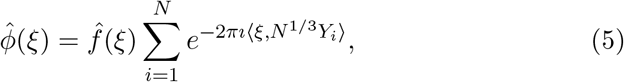

but now the CLT can no longer be applied in a straightforward manner, because the summands in (5) are random variables that depend on *N*.

### 2.2 Shape of container and decay rate of the Fourier transform

The representation (5) facilitates the calculation of any moment of 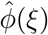. The expectation (first moment) of 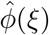 is given by

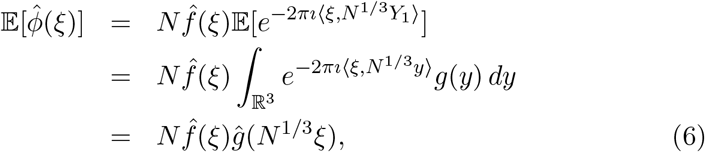

where *g*(*y*) is the probability density function of *Y*_1_, and *ĝ* is its Fourier transform. The dependency on *ĝ*(*N*^1/3^ξ) and *N* being a large parameter together suggest that the decay rate of *ĝ* at high frequencies is critical for analyzing Wilson statistics.

Different container shapes and choices of *g* can lead to different behavior of its Fourier transform *ĝ*. Before stating known theoretical results, it is instructive to consider a couple of examples.

- A uniform distribution in a ball. Here Ω_1_ is a ball of radius 1, denoted *B*, and the uniform density is 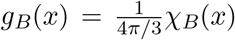, where *χ_B_* is the characteristic function of the ball. It is a radial function, a property that can readily be used to calculate its Fourier transform as

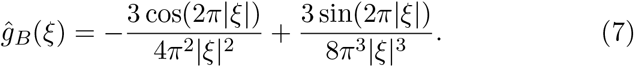 In particular, (7) implies that 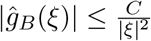 for some constant *C*.
- A uniform distribution in a cube. Here 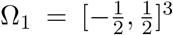 is the unit cube, and *g_C_* is a product of three rectangular window functions whose Fourier transform is the sinc function. As a result,

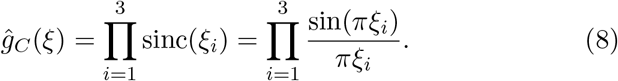 Taking ξ along one of the axes, e.g., ξ = (|ξ|,0,0) gives 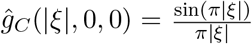. In this case, 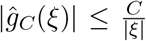 for some *C* > 0. Notice that the decay of *ĝ_C_* in directions not normal to its faces is faster. For example, for 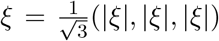 we have 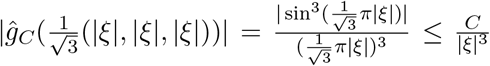.

We are now ready to state existing theoretical results about the decay rate of the Fourier transform for containers of general shape.

#### Theorem 1.

*(see [5], p. 336)*

1. *Suppose* 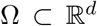 *is a bounded region whose boundary M* = *∂*Ω *has non-vanishing Gauss curvature at each point, then*

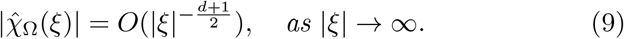
2. *If M has m non-vanishing principal curvatures at each point, then*

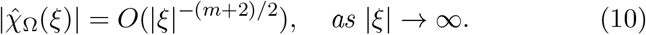 The decay rates previously observed for the three-dimensional ball (*d* = 3 or *m* = 2) and the cube (*m* = 0) are particular cases of Theorem 1. Although the decay rate in different directions could be different (as the example of the cube illustrates), for a large family of containers (convex sets and open sets with sufficiently smooth boundary surface), the following Theorem asserts that the spherical average of the power spectrum has the same decay rate as that of the ball.

#### Theorem 2.

*(see [6]) Suppose* 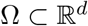 *is a convex body or an open bounded set whose boundary* ∂Ω *is C*^3/2^. *Then*,

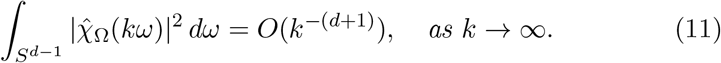

*Here k* = |ξ| *is the radial frequency and S*^*d*−1^ *is the unit sphere in* 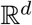.

### 2.3 Validity regime of Wilson statistics

We are now in position to state and prove our main result that fully characterizes the regime of validity of Wilson statistics.

#### Theorem 3.

1. *For the random bag of atoms model, the expected power spectrum is given by*

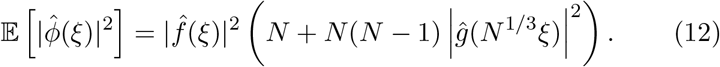
2. *If the container is a convex body or an open set with a C*^3/2^ *boundary surface, and the atom locations are uniformly distributed in the container, then the expected spherically-averaged power spectrum satisfies*

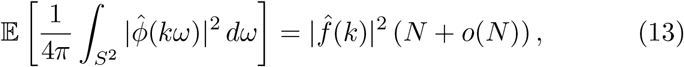

*for k* ≫ *N*^−1/12^.
3. *If the Fourier transform of the density g satisfies* |*ĝ*(ξ)| ≤ *C*|ξ|^−*α*^, *then*

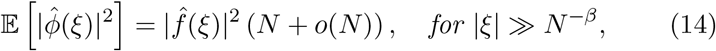

*where* 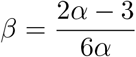.

*Proof*. Starting with Wilson’s original approach, from (5) it follows that the power spectrum of *ϕ* is given by

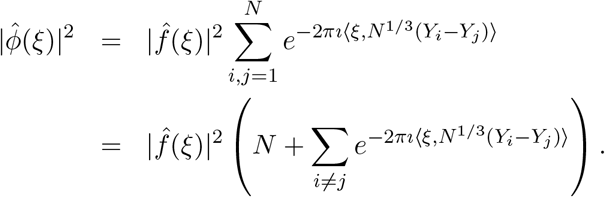

Since the *Y_i_*’s are i.i.d, the expected power spectrum satisfies

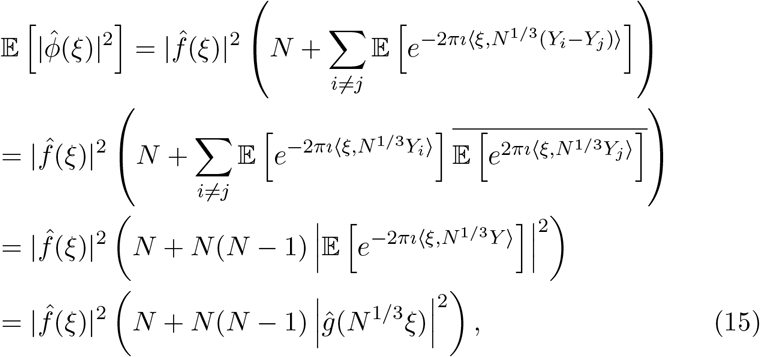

establishing (12). Assuming *f* (hence also 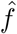) are radial functions, the expectation of the spherically-averaged power spectrum satisfies

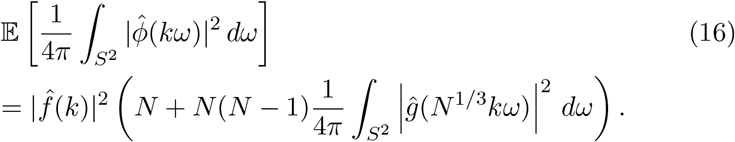

Theorem 2 with *d* = 3 implies

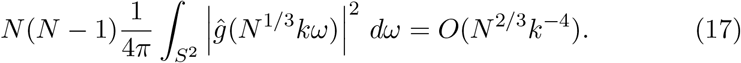

This term is negligible compared to *N* in (16) for *k* ≫ *N*^1/12^, proving (13). Finally, if |*ĝ*(ξ)| ≤ *C*|ξ|^−*α*^, then 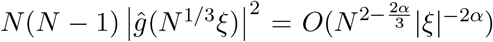, which is *o*(*N*) for 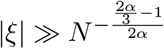 □

Note that (16) and (17) suggest that the spherically-averaged power spectrum decays to its high frequency limit as *k*^−4^. This decay rate at high frequencies is reminiscent of Porod’s law in SAXS [7, 8]. At first, the 1/12 exponent of the cutoff frequency *k*_0_ = *O*(*N*^−1/12^) might seem mysterious. In hindsight, it is simply the product of the dimension *d* = 3 that resulted in the scaling of *N*^1/3^ and the decay rate exponent of *k*^−4^.

### 2.4 Spherical averaging and statistical fluctuation

Note that in our derivation of Wilson statistics, we first took expectation with respect to the atom positions followed by spherically-averaging the power spectrum. On the other hand, spherically-averaging (3) first gives

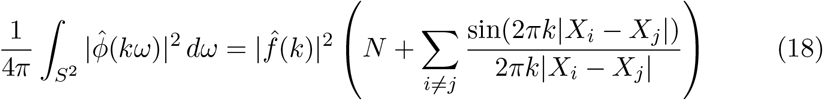

as in Debye’s scattering equation [9], due to the identity

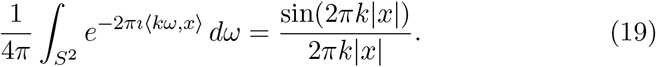

Although the 1/*k* decay of the sinc function in (18) sheds some light on the mechanism by which the sum over atom pairs decreases with *k*, it does not seem to provide a good starting point for a rigorous derivation of Wilson statistics, nor does it provide a clear path for the generalizations considered later in this paper.

While Theorem 3 characterizes the expected power spectrum, one may wonder whether the statistical fluctuations of the power spectrum could overwhelm its mean. This turns out not to be the case. Similar to the derivation of Wilson statistics, one can show that if |*ĝ*(ξ)| ≤ *C*|ξ|^−2^ then

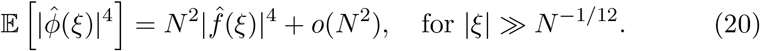

Since 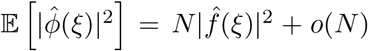 for |*ĝ*| ≫ *N*^−1/12^, it follows that for |*ĝ*| ≫ *N*^−1/12^

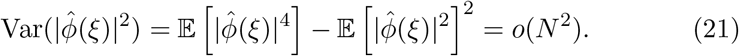

In other words, the standard deviation of the power spectrum is *o*(*N*), so the fluctuation is smaller than the mean value.

## 3 Theoretical Guinier plots and cutoff frequencies

A realistic estimate of the density of atoms in proteins gives rise to theoretical Guinier plots and prediction of the cutoff frequency above which Wilson statistics holds. The protein density is approximately *ρ* ≈ 0.8Da/Å^3^ [10]. The number of carbon atom equivalents, using 9.1 carbon equivalents per amino acid of molecular weight 110 is 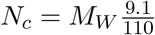, where *M_W_* is the molecular weight. For a spherically-shaped protein of radius *R*, the molecular weight and number of carbon atom equivalents are given by 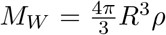 and 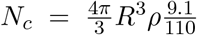, respectively. In particular, *N_c_* = 2.77 × 10^5^ and *M_W_* = 3.3MDa for *R* = 100Å, while *N_c_* = 4.3 × 10^3^ and *M_W_* = 52kDa for *R* = 25Å (see Table 2 in [10]).

**Figure 1:**
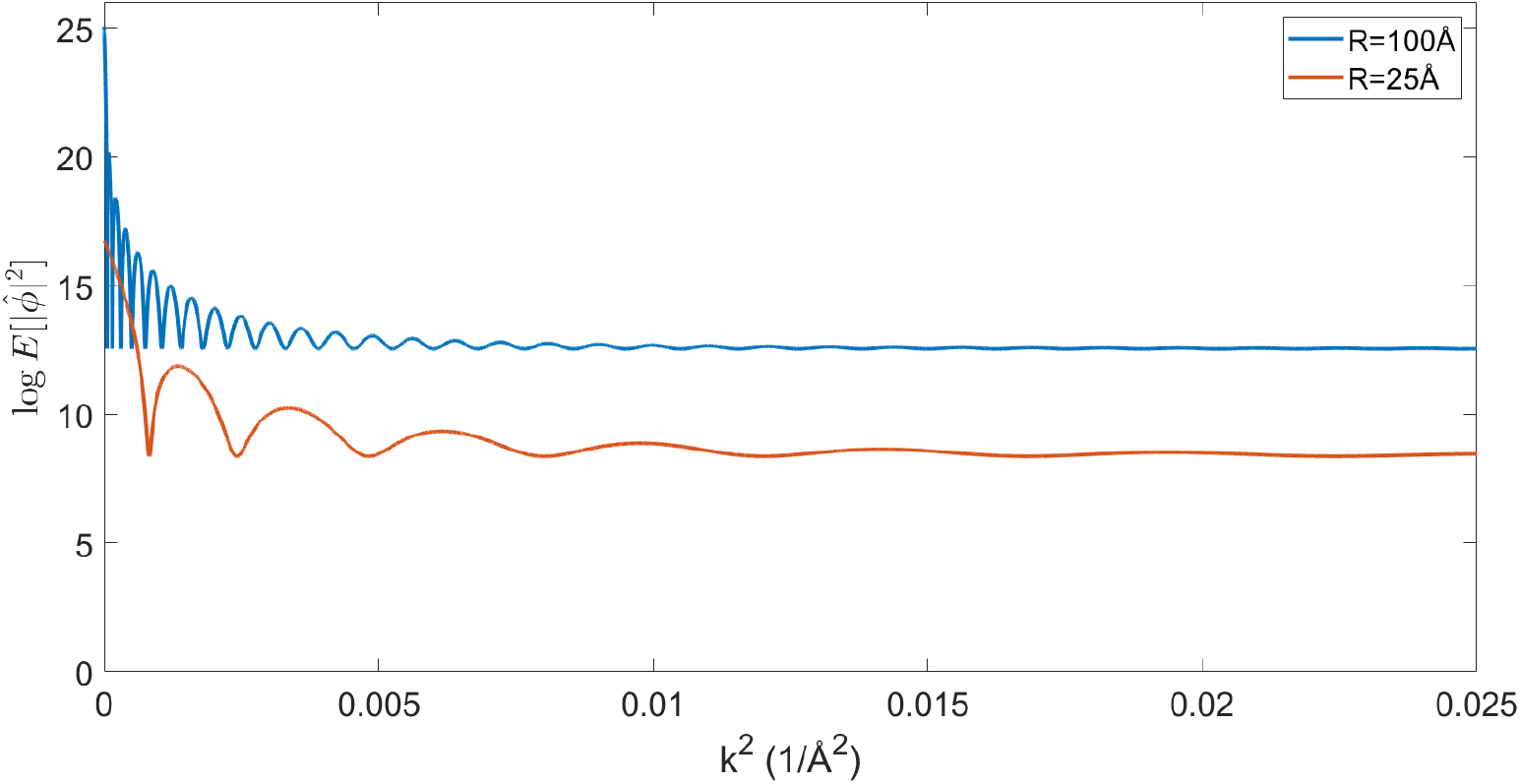
Theoretical Guinier plots as predicted by Theorem 3 for realistic uniform density of atoms in balls of radius 25Å and 100Å.

**Figure 2:**
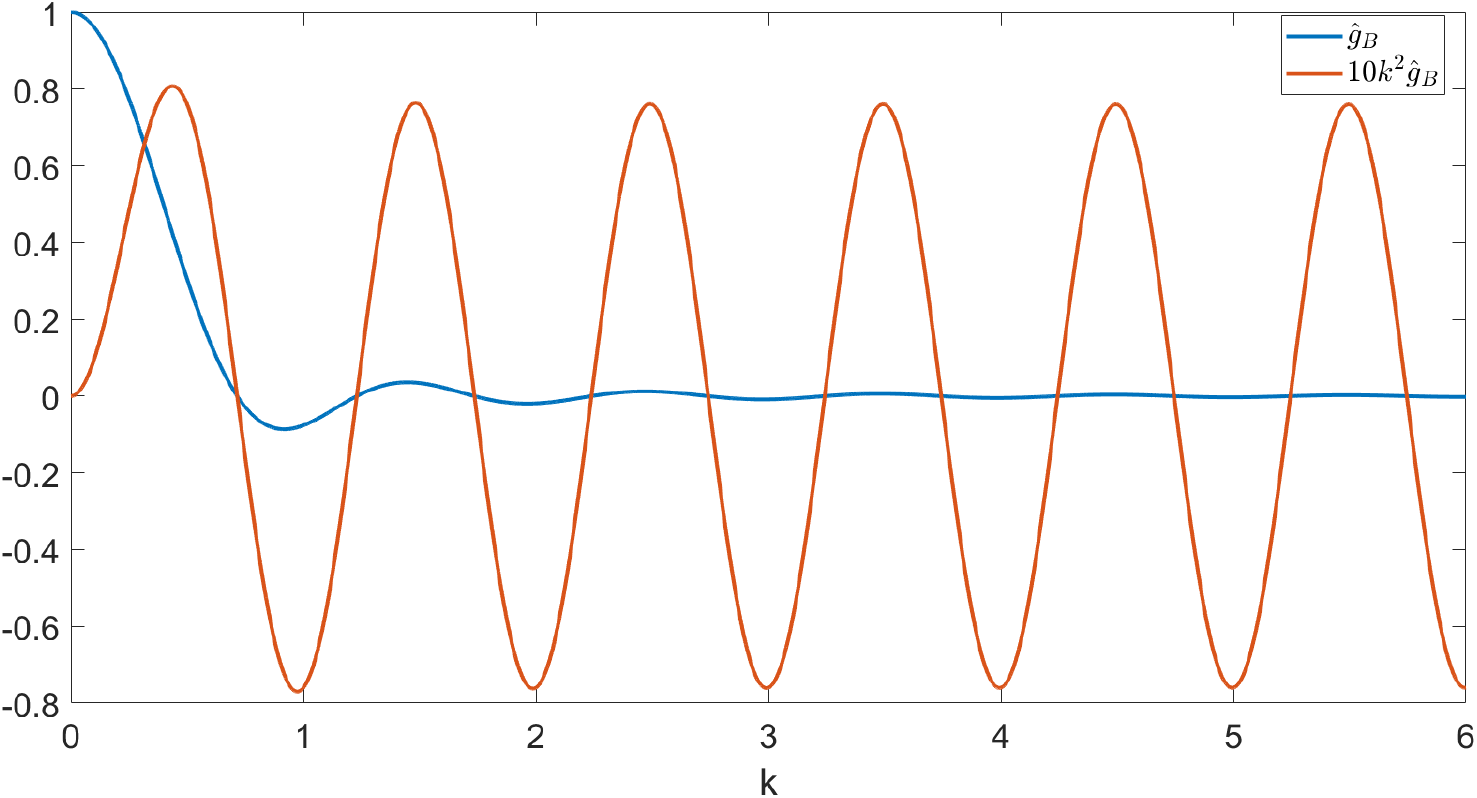
A closer look at the Fourier transform of the uniform density in the unit ball *ĝ_B_* given by (7). The radial frequency *k* = |ξ| is dimensionless here.

Theoretical Guinier plots of the logarithm of the expected power spectra (using (12) and (7)) as a function of the squared spatial frequency for these representative cases are shown in Figure 3. The effect of the atomic structure factor is not included in Figure 3 for which 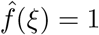. Also not included is the modification due to solvent contrast. The low-frequency signal is modified by the partial contrast-matching of solvent. In [2] the remaining contrast is estimated to be 0.42, so the low-frequency spectral density should be modified by this.

**Figure 3:**
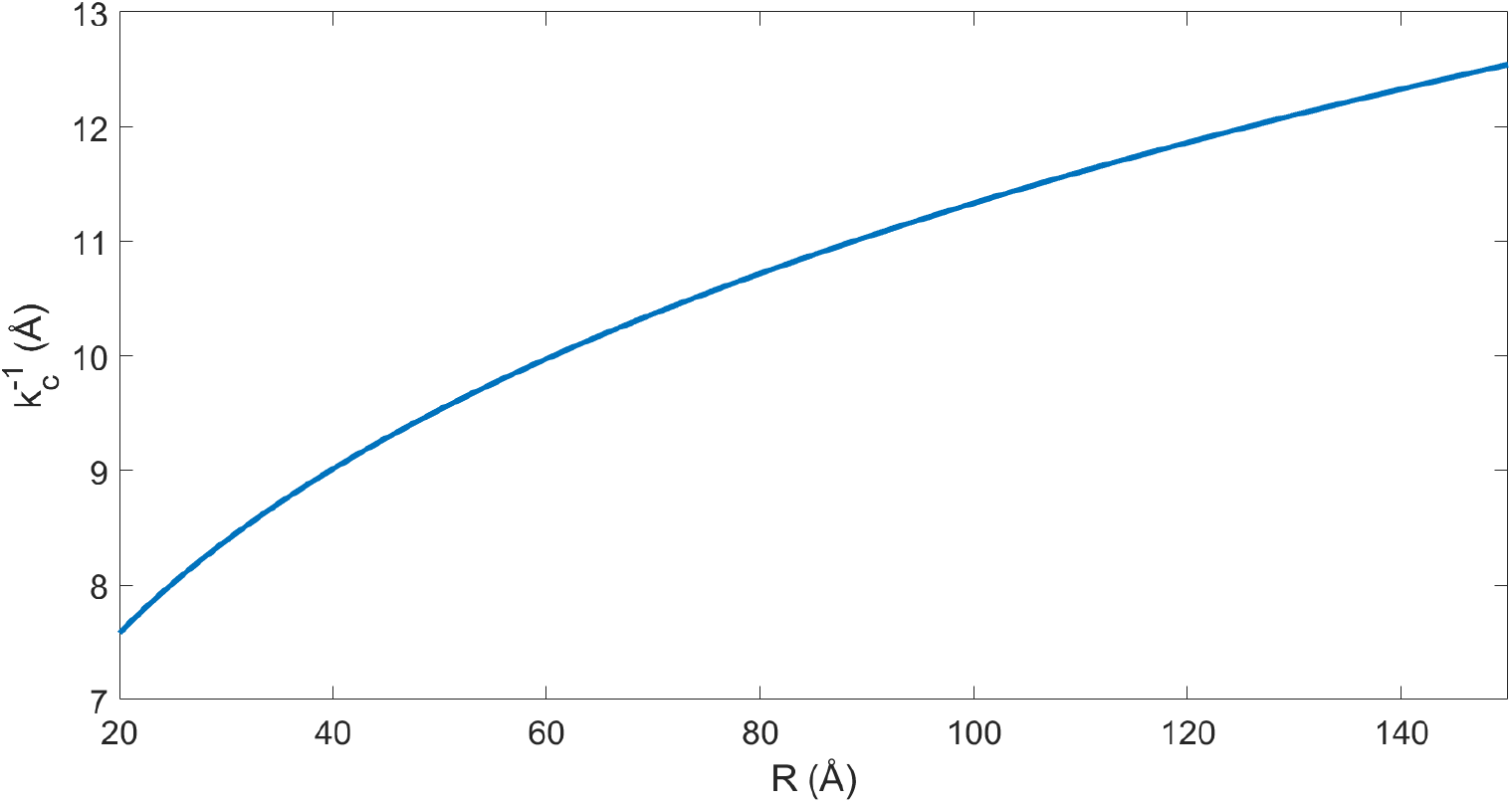
The cutoff resolution 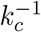 as a function of the radius *R* of a spherical protein with uniform distribution of atoms as given by (22).

The theoretical Guinier plots qualitatively resemble experimental Guinier plots, such as Figure 8 in [2]. For the larger molecule with *R* = 100Å the power spectrum is approximately flat above *k*^2^ = 0.01Å^−2^ corresponding to 10Å resolution, whereas for the smaller molecule with *R* = 25Å the transition occurs closer to *k*^2^ = 0.015Å^−2^, or 8.2Å resolution.

The notable oscillations in the Guinier plots are due to the oscillations of *ĝ_B_* given by (7). Figure 3 shows *ĝ_B_* and *k*^2^*ĝ_B_* (the latter is multiplied by 10 in order to make the two plots comparable in scale). We see that 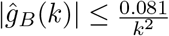 (i.e., the constant *C* in 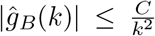 can be taken as *C* = 0.081).

It is important to keep in mind that proteins are not perfectly spherically symmetric. Although oscillations in the Guinier plot are still expected (and are indeed observed), their magnitude and periodicity are shape dependent.

Theorem 3 implies that the transition to Wilson statistics in the Guinier plot occurs at 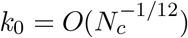, and for higher radial frequencies the spherically-averaged power spectrum is approximately flat. The cutoff frequency can be determined by balancing the two terms in (12). Specifically, we require the second term of (12) to be at most 0.3*N_c_*. This criterion, together with the bound 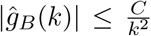 with *C* = 0.081 imply 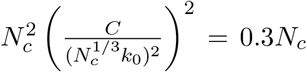, or 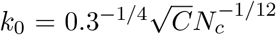. The radius *R*_0_ of the unit cell (that occupies a single atom on average) satisfies 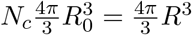. Therefore, 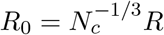, and the dimensional cutoff frequency *k_c_* (in Å^−1^) is given by

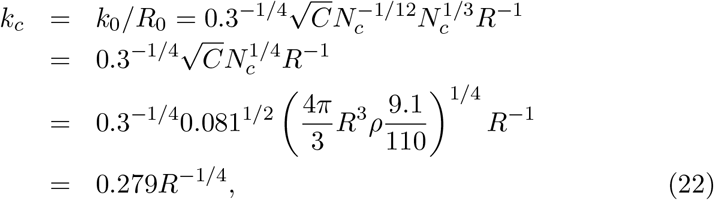

in terms of the radius, or equivalently

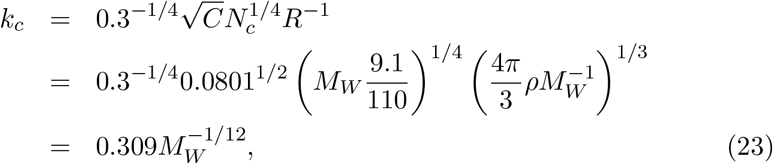

in terms of the molecular weight. The cutoff frequency decreases with the size of the molecule, but the decrease is quite gradual due the small exponent 1/12 in (23). For example, the cutoff frequency increases by just 47% when the molecular weight decreases by a factor of 100. For a large macromolecule with *M_W_* = 3.3MDa and *R* = 100Å the cutoff frequency is *k_c_* = 0.088Å^−1^ corresponding to 11.3Å resolution. For a smaller macromolecule with *M*_W_ = 52kDa and *R* = 25Å the cutoff frequency is *k_c_* = 0.125Å^−1^ corresponding to 8.0Å resolution. These predictions are in agreement with our previous estimates for the cutoff frequencies that were obtained by eyeballing Figure 3. Figure 3 illustrates the cutoff frequency as a function of the molecular size with radius extremes of 20Å to 150Å. The cutoff frequency is relatively stable and varies only little across a wide range of molecular sizes (from 7.5Å to 12.5Å resolution). This behavior and resolutions are in agreement with empirical evidence about the validity regime of Wilson statistics [2].

## 4 Generalizations and applications to cryo-EM

### 4.1 Existing applications to cryo-EM

A common practice in single particle cryo-EM is to apply a filter to the reconstructed map. The filter boosts medium and high frequencies such that the power spectrum of the sharpened map is approximately flat and consistent with Wilson statistics [2, 11]. The filter is an exponentially growing filter whose parameter is estimated using the Guinier plot. The boost of medium and high frequency components increases the contrast of many structural features of the map and helps to model the atomic structure. This is the so-called B-factor correction, B-factor flattening, or B-factor sharpening. It is a tremendously effective method to increase the interpretability of the reconstructed map. In fact, most map depositions in the Electron Microscopy Data Base (EMDB) only contain sharpened maps [12]. Map sharpening is still an active area of research and method development, see, e.g. [13, 14] and references therein. Wilson statistics is also used to reason about and extrapolate the number of particles required to high resolution [2].

### 4.2 Generalization of Wilson statistics to covariance with application to 3-D iterative refinement

We now highlight a certain generalization of Wilson statistics with potential application to 3-D iterative refinement, arguably the main component of the computational pipeline for single particle analysis [15]. Specifically, the Bayesian inference framework underlying the popular software toolbox RELION [16] requires the covariance matrix of 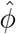 and approximates it with a diagonal matrix [17]. For tractable computation, the variance (the diagonal of the covariance matrix) is further assumed to be a radial function.

The random bag of atoms model underlying Wilson statistics provides the covariance matrix

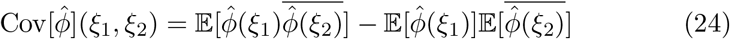

in closed form as

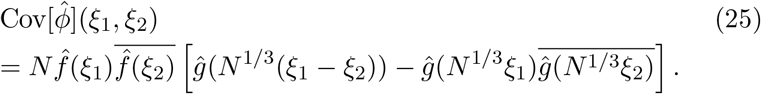

Before proving this result, note that it implies a vast reduction in the number of parameters needed to describe the covariance matrix. In general, for a 3-D map represented as an array of *L*^3^ voxels, the covariance matrix is of size *L*^3^ × *L*^3^ which requires *O*(*L*^6^) entries, which is prohibitively large. However, (25) suggests that the covariance depends on only *O*(*L*^3^) parameters. Furthermore, approximating *ĝ*(ξ) by a radial function implies that the covariance depends on just *O*(*L*) parameters, the same number of parameters in the existing Bayesian inference method for 3-D iterative refinement. Moreover, comparing the two terms in (25), the decay of *ĝ* implies that 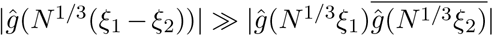 whenever |ξ_1_|,|ξ_2_| ≫ *N*^−1/3^. Therefore, for |ξ_1_|,|ξ_2_| ≫ *N*^−1/3^

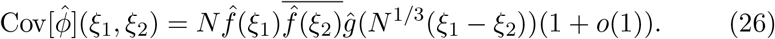

Since *ĝ*(*N*^1/3^(*ĝ*_1_–*ĝ*_2_)) is largest for ξ_1_ = ξ_2_ and decays with increasing distance |ξ_1_ – ξ_2_|, it follows from (26) that the covariance matrix restricted to frequencies above *N*^−1/3^ is approximately a band matrix with bandwidth *O*(*N*^−1/3^), such that the diagonal is dominant and matrix entries decay when moving away from the diagonal. Note that *N*^−1/3^ is a very low frequency corresponding to resolution of the size of the protein (as implied by the *N*^1/3^ scaling). Therefore, the covariance is well approximated by a band matrix with a very small number of diagonals. This serves as a theoretical justification for the diagonal approximation in the Bayesian inference framework [17], as correlations of Fourier coefficients with |ξ_1_ –ξ_2_| ≫ *N*^−1/3^ are negligible. On the flip side, correlations for which |ξ_1_ – ξ_2_| ≪ *N*^−1/3^ should not be ignored and correctly accounting for them could potentially lead to further improvement of [17].

To prove (25), we evaluate the two terms in the right hand side of (24) separately. The second term is directly obtained from (6) as

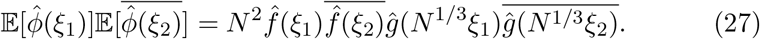

To evaluate the first term, we substitute 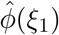 and 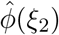 by (5), separate the summation into diagonal terms (*i* = *j*) and off-diagonal terms (*i* ≠ *j*) as in Wilson’s original argument, and use that *Y_i_*’s are i.i.d, resulting

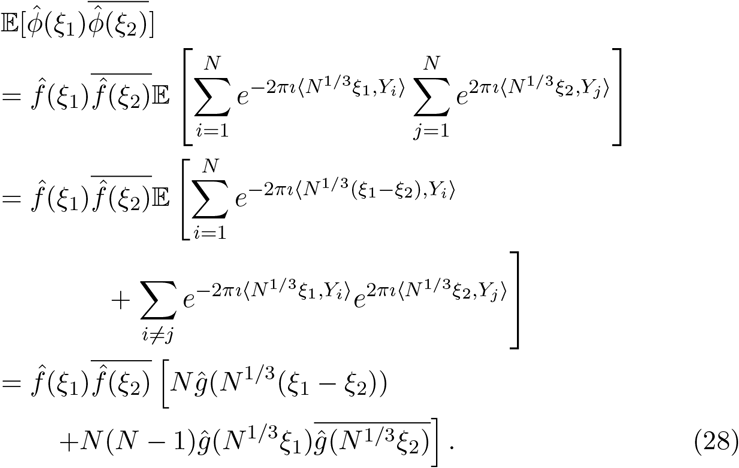

Subtracting (27) from (28) proves (25). This is a generalization of Wilson statistics, as setting ξ_1_ = ξ_2_ reduces (28) to (12).

Note that the diagonal of the covariance matrix satisfies

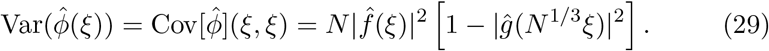

The variance vanishes for ξ = 0 because 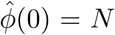 regardless of the atoms positions. The small variance at very low frequencies shares the same origins of Guinier law.

In existing Bayesian inference approaches [17], the mean of each frequency voxel is assumed to be zero. However, comparing (6) and (29) for the mean 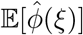 and the variance 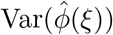, we see that the variance dominates the squared mean only for |ξ| ≫ *N*^−1/12^, which is the validity regime of Wilson statistics. It follows that it is justified to assume a zero-mean signal only for high frequencies, but not at low frequencies. Including an explicit (approximately radial) non-zero mean in the Bayesian inference framework may therefore bring further improvement.

### 4.3 Generalization of Wilson statistics to higher order spectra with application to autocorrelation analysis

Autocorrelation analysis, originally proposed by Kam [18, 19], has recently found revived interest for experiments using X-ray free electron laser (XFEL) [20, 21, 22] and cryo-EM [23, 24, 25]. In autocorrelation analysis, the three-dimensional molecular structure is determined from the correlation statistics of the noisy images. Typically, the second or third order correlation functions are sufficient in principle to uniquely determine the structure [26, 23]. It is therefore of interest to derive a third order statistics analogue of (12). Specifically, 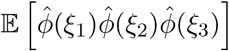 is given by

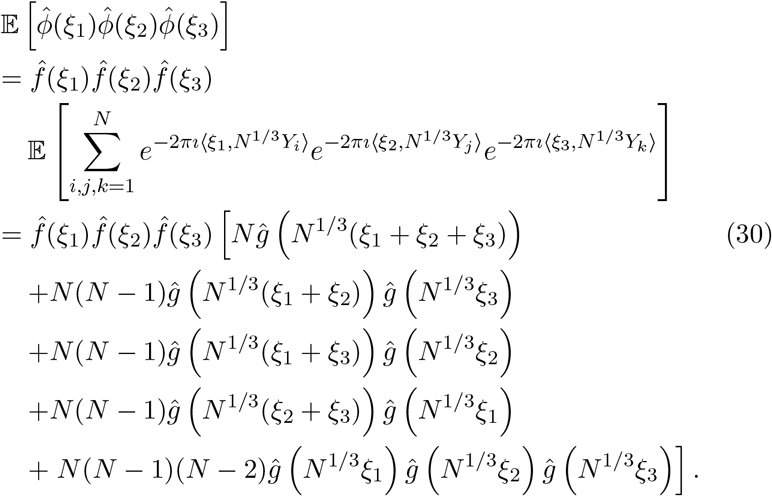

This result is obtained by separating the sum over all triplets *i, j, k* into five groups: *i* = *j* = *k*, *i* = *j* ≠ *k*, *i* = *k* ≠ *j*, *j* = *k* ≠ *i*, and *i* ≠ *j* ≠ *k* = *i*.

Similar to the power spectrum 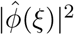 which is the Fourier transform of the autocorrelation function, the bispectrum 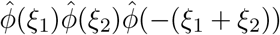 is the Fourier transform of the triple-correlation function. The bispectrum, like the power spectrum, is also shift-invariant. As such, it plays an important role in various autocorrelation analysis techniques. The expected bispectrum under the random bag of atoms model is obtained by setting ξ_1_+ξ_2_+ξ_3_ = 0 in (31)

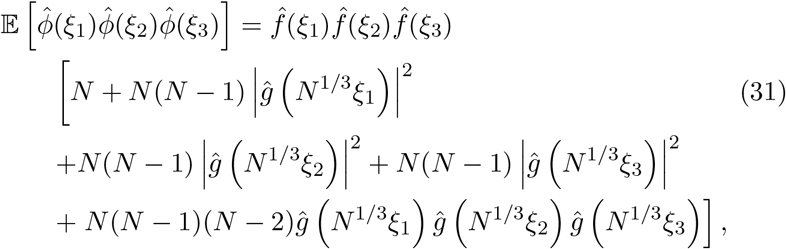

for ξ_1_ + ξ_2_ + ξ_3_ = 0.

The bispectrum drops from *N*^3^ for ξ_1_ = ξ_2_ = ξ_3_ = 0 to *N* at high frequencies. This drop is even more pronounced than that of the power spectrum that decreases from *N*^2^ to *N*. This may lead to numerical difficulties in inverting the bispectrum as it has a large dynamic range, e.g., it spans eight orders of magnitude for *N* = 10^4^.

The terms in the first two lines of (31) have similar behavior to the power spectrum (12). The last term depends on the decay rate of *ĝ*. If |*ĝ*(ξ)|≤*C*|ξ|^−2^ as for the ball, then

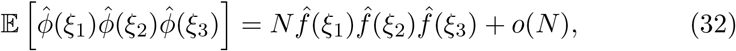

for |ξ_1_|,|ξ_2_|,ξ|_3_| ≫ 1, which can be regarded as a generalization of Wilson statistics (e.g., (13)) to higher order spectra. However, for higher order spectra such as the bispectrum the behavior at high frequencies is more involved. For example, taking ξ_1_ and ξ_2_ to be high frequencies does not imply ξ_3_ = –(ξ_1_ + ξ_2_) is necessarily a high frequency, as can be readily seen by taking ξ_2_ = −ξ_1_ for which ξ_3_ = 0. For this particular choice of ξ_2_ = −ξ_1_ the expected bispectrum is always greater than *N*^2^.

## 5 Discussion

This paper provided the first formal mathematical derivation of Wilson statistics, offered generalizations to other statistics, and highlighted potential applications in structural biology.

The assumption underlying Wilson statistics of independent atom locations is too simplistic as it ignores correlations between atom positions in the protein. It is well known that the power spectrum deviates from Wilson statistics at frequencies that correspond to interatomic distances associated with secondary structure such as *α*-helices which produce a peak at 10Å and beta-sheets which produce a peak at 4.5Å. A more refined model that includes such correlations is beyond the scope of this paper.

From the computational perspective, we note that numerical evaluation of Fourier transforms and power spectra associated with Wilson statistics involves computing sums of complex exponentials of the form (1). These can be efficiently computed as a Type-1 3-D non-uniform fast Fourier transform (NUFFT) [27]. The computational complexity of a naïve procedure is *O*(*NM*), where *M* is the number of target frequencies, whereas the asymptotic complexity of NUFFT is *O*(*N* + *M*) (up to logarithmic factors). These considerations will be taken into account in future computational work for numerical validation of the theoretical predictions including comparison with the power spectra and bispectra of density maps created from atomic models [28].

Wilson statistics is an instance of a universality phenomenon: all proteins regardless of their shape and specific atomic positions exhibit a similar spherically-averaged power spectrum at high frequencies. From the computational standpoint in cryo-EM, this universality is a blessing and a curse at the same time. On the one hand, it enables to correct the magnitudes of the Fourier coefficients of the reconstructed map so they agree with the theoretical prediction. On the other hand, it implies that the high frequency part of the spherically-averaged power spectrum is not particularly useful for structure determination, as it does not discriminate between molecules. The generalization of Wilson statistics to the higher order spectra shows that the bispectrum also becomes flat at high frequencies. These observations may help explain difficulties of the autocorrelation approach as a high resolution reconstruction method [24].

## Acknowledgements

The author is indebt to Nicholas Marshall, Fred Sigworth, Ti-Yen Lan, Tamir Bendory, and Joe Kileel for valuable discussions and comments. This work was supported in part by AFOSR Awards FA9550-17-1-0291 and FA9550-20-1-0266, the Simons Foundation Math+X Investigator Award, the Moore Foundation Data-Driven Discovery Investigator Award, NSF BIGDATA Award IIS1837992, NSF Award DMS-2009753, and NIH/NIGMS Award R01GM136780-01.

